# Socioeconomic predictors of knockdown resistance in *Aedes albopictus* (Diptera: Culicidae)

**DOI:** 10.1101/2025.07.01.662680

**Authors:** Cole D. Butler, Jessica Y. Ding, Jennifer Baltzegar, Zachary S. Brown, Martha O. Burford Reiskind, Michael H. Reiskind

## Abstract

Knockdown resistance (*kdr*) in the mosquito *Aedes albopictus* (Skuse) jeopardizes the effectiveness of insecticidal control. This is a pressing issue given the expanding range of the species and its role as vector to harmful viruses. Effectively preventing the emergence of resistance or removing the conditions that positively select for *kdr* mutations requires us first to understand how these conditions arise. Here, we investigate the association between wealth and the frequency of *kdr* in *Ae. albopictus* populations in Raleigh, North Carolina, USA. We hypothesized that *kdr* frequency correlates with wealth, measured by total property value. We speculate that wealthier neighborhoods apply chemical insecticides more frequently, leading to higher *kdr* frequencies. We tested this hypothesis by sampling mosquito populations from 31 different residential blocks across the city and along a property value gradient. We found a high frequency of 39.0% for mutations at locus F1534S of the voltage-gated sodium channel gene (*vgsc*). *Kdr* mutations were found at 84% of the blocks we sampled. Our statistical analysis indicates strong evidence for an association between wealth and F1534S frequency. We discuss these and other findings, and what this means for suburban mosquito control going forward.

## 1 Introduction

The tiger mosquito *Aedes albopictus* (Skuse) is an invasive pest with a global distribution, including the East Coast of the United States from Florida to Connecticut [2, 9, 35], and recent reports as far north as Massachusetts and New Hampshire [27]. First introduced to the continental United States in Texas in the mid-1980s [46], *Ae. albopictus* quickly spread to North Carolina where, by 1998, it was reported in all counties [45]. Today, *Ae. albopictus* is the most common mosquito species in the state and is a nuisance pest in suburban yards [61, 54]. Aedes albopictus is peridomestic and closely associated with humans in both its feeding and oviposition habits. The tiger mosquito commonly feeds on humans but is known to take bloodmeals from other animals, including dogs and cats [17, 21, 57]. Females prefer to oviposit in artificial water containers such as plastic bottles, cans, and pots that are commonly found around human dwellings [11, 53].

Beyond it being an aggressive biter and a major nuisance, *Ae. albopictus* is a public health concern for its established role in transmitting the pathogens responsible for dengue, West Nile, and chikungunya. Dengue alone infected over 7 million people in the Americas in 2024, and case counts continue to increase [75]. *Ae. albopictus* has been heavily implicated in dengue outbreaks in non-endemic regions [16, 32, 49], and in the United States has played pivotal roles in the Hawaii dengue epidemics in 2001-2002 [16] and again in 2015-2016 [29]. Indeed, previous research has suggested that it is possible for *Ae. albopictus* to sustain local transmission of dengue and other arboviruses in the continental United States [25]. Wild-caught *Ae. albopictus* samples have also been found carrying strains of other arboviruses, including Eastern equine encephalitis [10] and La Crosse viruses [43]. *Ae. albopictus* is also responsible for transmitting the filarial parasite that causes heartworm disease in dogs [36].

North Carolina lacks state-funded mosquito control resources and Wake County similarly does not offer mosquito control services [55]. This means that all mosquito control in Raleigh, NC is performed at the household level by residents themselves or contracted out to private companies. Suburban mosquito control primarily relies on the spraying of insecticides, with pyrethroids being among the most common. Pyrethroids disrupt normal function by binding to transmembrane proteins in nerve cells, causing a knockdown effect. Nonsynonymous changes in the voltage-gated sodium channel (*vgsc*) gene shift amino acids and alter the transmembrane protein, reducing insecticide sensitivity and resulting in knockdown resistance (*kdr*). These changes confer resistance to both dichlorodiphenyltrichloroethane (DDT) and pyrethroids. The most common target site mutation associated with *kdr* in *Ae. albopictus* is at locus F1534S of the *vgsc* gene.

The first report of *kdr* in *Ae. albopictus* was in Singapore in 2009 [33]. Since then, *kdr* mutations have been reported in *Ae. albopictus* samples all over the world, including China [77], Spain [6], Italy [50], and the United States [38, 40] (see reviews in [3, 65] for more details). In the United States, phenotypic resistance to specific pyrethroids has been observed in *Ae. albopictus* as early as 2004 in Alabama and Florida [38]. In 2011, Marcombe *et al*. (2014) reported phenotypic resistance to DDT in mosquitoes sampled from Florida and New Jersey. Genetic sequencing revealed *kdr* mutations at locus 1534 in domain III of the *vgsc* gene. *Kdr* mutations in North Carolina populations of *Ae. albopictus* are a recent phenomenon. A 2014 study failed to detect phenotypic resistance to bifenthrin (a Type I pyrethroid) [67], while a 2017 survey of Mecklenburg County failed to find any genetic indicators of *kdr*, although they did not sequence domain III of the *vgsc* gene [48]. Also in 2017, Richards *et al*. (2019) collected eggs from seven counties across North Carolina, including Wake and Mecklenburg counties [58]. The investigators found that some populations were resistant or developing resistance to the pyrethroids permethrin and tau-fluvalinate. In 2020, investigators found 12 *Ae. albopictus* mosquitoes in Wake County (out of 412) with a mutation at locus 1534 that substitutes serine (S) for the wild-type phenylalanine (F); no other *kdr* mutations were found [1]. This specific mutation, F1534S, has consistently been shown to correlate with pyrethroid resistance in *Ae. albopictus* [12, 23, 62, 76]. Using *Ae. albopictus* samples collected in Raleigh, NC as early as 2016, we found in a separate study that the Ser1534 mutation first appeared in 2018 [5]. The 2023 dataset analyzed in this work is part of a larger temporal trend reported in Baltzegar *et al*. (2025).

Noting the restricted distribution of resistant alleles in Wake County in 2020 to one wealthy, older neighborhood, we hypothesized that the frequency of *kdr* in *Ae. albopictus* is correlated with wealth and mosquito abundance. We tested our hypothesis by predicting that wealthier neighborhoods (where people can afford private mosquito control) and older neighborhoods (where previous data show higher mosquito abundance [61]) will have a higher frequency of F1534S alleles than poorer, newer neighborhoods. To assess our predictions, we conducted a survey of 31 grid blocks for mosquito abundance and F1534S frequency across a neighborhood wealth and age gradient.

## 2 Methods

### 2.1 Study Area

Collections took place in Raleigh, Wake County, NC. Raleigh is a city of nearly half a million people that includes densely urban blocks and sprawling suburban neighborhoods [69]. Socioeconomic disparities in Raleigh are stark: the median household income of census blocks ranges from below $30,000 to over $250,000 [71]. 11.4% of the population lives in poverty [66]. We exclusively sampled from neighborhoods zoned as residential—either four or six homes per acre (designated R-4 and R-6, respectively). All sampling sites were single-family homes with lawns.

### 2.2 Site Selection

We selected 40 blocks at random from a grid map of all R-4 and R-6 areas in Raleigh, where each block has an area of 1 km^2^. Block size was chosen so that we could assume that the mosquito populations in separate blocks are (approximately) genetically independent, based on the dispersal habits of *Ae. albopictus*. The average dispersal of *Ae. albopictus* over its lifetime is estimated to be 200-400 m. [42, 63]. While dispersal over longer distances has been reported (e.g., [31]), it is typical that most mosquitoes will disperse less than 1 km. from their natal site [42]. Dispersal is likely further reduced given the abundance of oviposition sites and hosts in suburban neighborhoods [56], and the presence of busy roads. Any blocks with less than 50% of its area zoned as R-4 or R-6 were discarded. Within each block, sampling took place at houses located near the block centroid. If the block centroid did not fit our selection criteria (e.g., an apartment complex), the block was discarded. This left a total of 31 blocks (see Fig. S1). Save for one instance (Block 16), all of the houses within each block were located on the same street.

### 2.3 Mosquito Sampling

We collected mosquitoes over a one-month period from mid-August to mid-September 2023 using BG-Sentinel traps baited with scent lures and dry ice. One trap was used per sampling site and was left for approximately 24 hours before collection. Sites at which the number of collected mosquitoes was low were re-sampled to improve our estimated average *kdr* frequency per block. In instances where homeowners were not receptive to re-sampling, a different house was chosen nearby. Of the 40 houses at which we conducted sampling, 22 houses were sampled twice, and two houses were sampled three times. Mosquitoes were identified by morphological characteristics and a subset of those collected—up to 32 samples per block— was chosen for DNA extraction and sequencing.

### 2.4 DNA Extraction and Sequencing

Genomic DNA (gDNA) was extracted using a Qiagen DNeasy Kit (catalog number 69506) following the manufacturer’s protocol with the following modifications: homogenized whole mosquitoes were incubated at 55*°*C with 180 ml ATL buffer and 20*µ*l Proteinase K overnight. An RNase A treatment was performed at the beginning of the day 2 extraction. Purifying washes were then completed as prescribed and gDNA was eluted in 100ul dH2O for downstream analyses.

### 2.5 Genotyping *kdr* Mutation

Allele-specific polymerase chain reaction (AS-PCR) and melting curve analysis was performed on gDNA from individual mosquitoes to determine the genotype at *kdr* locus F1534S using a Bio-Rad CFX384 Touch Real-Time PCR Detection System (Hercules, CA) in the Genomic Sciences Laboratory at North Carolina State University according to the protocol by Baltzegar *et al*. (2025). A total of three primers were used in this protocol including 1 common reverse primer (F1534R: TCT GCT CGT TGA AGT TGT CGA T), and 2 forward primers—one for each of the potential alleles. Primer S1534F (GCG GGC AGG GCG GCG GGG GCG GGG CCT CTA CTT CGT GTT CTT CAT CAT GTC) is designed to amplify the resistance allele and primer F1534F (GCG GGC TCT ACT TCG TGT TCT TCA TCA TAT T) is designed to amplify the susceptible allele. The PCR volume for each reaction was 10 *µ*l and included 5*µ*l dH2O, 0.2 *µ*l S1534F, 0.4 *µ*l F1534F, 0.4 *µ*l F1534R, 3.0 *µ*l PerfeCTa SYBR Green SuperMix for iQ (VWR – 101414-144), and 1.0 *µ*l of gDNA. Thermal cycling conditions were: step 1—95^*°*^C for 2 min, step 2—95^*°*^C for 30 s, step 3—58^*°*^C for 30 s, step 4—72^*°*^C for 30 s, step 5—go to step 2, 34 times, step 6—72^*°*^C for 2 min, and step 7—melting curve 65–95^*°*^C, increment 0.2^*°*^C per 10 s plus a plate read. Analysis of melting curve peaks was performed on CFX Maestro Software (Bio-Rad, Hercules, CA) and verified by eye. Each sample was genotyped twice to ensure accuracy of results. Samples were discarded from further analysis if results varied between the two runs.

### 2.6 Data

We used property value as a proxy for wealth. The property value of each collection site was retrieved from Wake County’s Computer Assisted Mass Appraisal system, as reported on the Residential Sales Search tool [70]. Property value includes the value of the house and the lot. In addition to property value, we used house age and abundance as explanatory variables. House age is included based on findings that *Ae. albopictus* abundance increases with neighborhood age [61] and might thus affect the frequency of insecticide spraying. Mosquito abundance is the total number of mosquitoes caught per 24-hour trapping period. When more than one house was sampled in a given block, we averaged all data across houses to produce one data point per block. We log-transformed total property values to reduce the effect of outliers and mean-centered all predictors before statistical analysis.

## 3 Statistical Analysis

We take as our response variable the allele frequency of the *kdr* mutation Ser1534 for each block (*n* = 31). We first used Moran’s *I* to test for spatial autocorrelation of *kdr* allele frequencies [24]. Moran’s *I* is a global measure of spatial autocorrelation and ranges from –1 to 1, with 0 representing no spatial autocorrelation, *I <* 0 representing negative autocorrelation, and *I >* 0 representing positive autocorrelation [18]. We then investigated the relationship between *kdr* allele frequency and our explanatory variables with and without an interaction term between house age and mosquito abundance. In both cases, we fit a logistic model with the canonical link function in which resistant alleles are considered “successes” and treat block as a random variable. After this, we used the quasi-Akaike Information Criterion corrected for small sample sizes (QAICc) to select the most parsimonious multivariate model [8]. To check that the models were properly specified, we used Moran’s *I* to test for spatial independence of the residuals [24]. Model residuals exhibited no evidence of dependence or spatial autocorrelation based on Moran’s *I*. This was regardless of whether we assumed neighboring blocks share an edge (so-called “rook” contiguity) or share an edge/vertex (“queen” contiguity). We also investigated mosquito abundance per sampling site (*n* = 40) with house age and total property value as predictors, based on findings from a previous study [61]. To do this, we fit a negative binomial model with the number of 24-hour trapping periods as an offset. As found in *kdr* frequency, an analysis of model residuals indicated no evidence of dependence nor spatial autocorrelation based on Moran’s *I* using an inverse distance matrix. For both models, we use a non-parametric bootstrap to conduct hypothesis tests [28] and compute confidence intervals for the expected response and regression coefficients. All analysis was done in R 4.4.3 [52]; the negative binomial model was fitted using the glm.nb function in the MASS package [68].

## 4 Results

The blocks included in our study spanned a large gradient of average house age (by build year, min = 1940; Q1 = 1963; median = 1971; Q3 = 1988; max = 2022) and total property value (min = $183,320; Q1 = $370,605; median = $523,507; Q3 = $778,361; max = $2,783,555). We note that house age and total property value were not significantly correlated in our dataset (*ρ* = 0.07). We captured 987 *Ae. albopictus* mosquitoes, with an average of 19.5 per trap. Trap counts were highly skewed and ranged from 0-200 mosquitoes (Fig. S1). Of 583 mosquito samples chosen for AS-PCR, 574 were genotyped successfully and 535 produced consistent results when genotyped a second time. The Ser1534 frequency per block was 37.7%, on average. The only blocks for which we did not find mosquitoes with *kdr* mutations were all located in the southeastern part of the city (Fig. 1). The overall Ser1534 frequency (pooling all samples) was 39.0%. A breakdown of pooled genotype frequencies is as follows: homozygous susceptible (S/S; 47.9%), heterozygous (R/S; 26.4%), and homozygous resistant (R/R; 25.8%) (Fig. 1).

**Figure 1.**
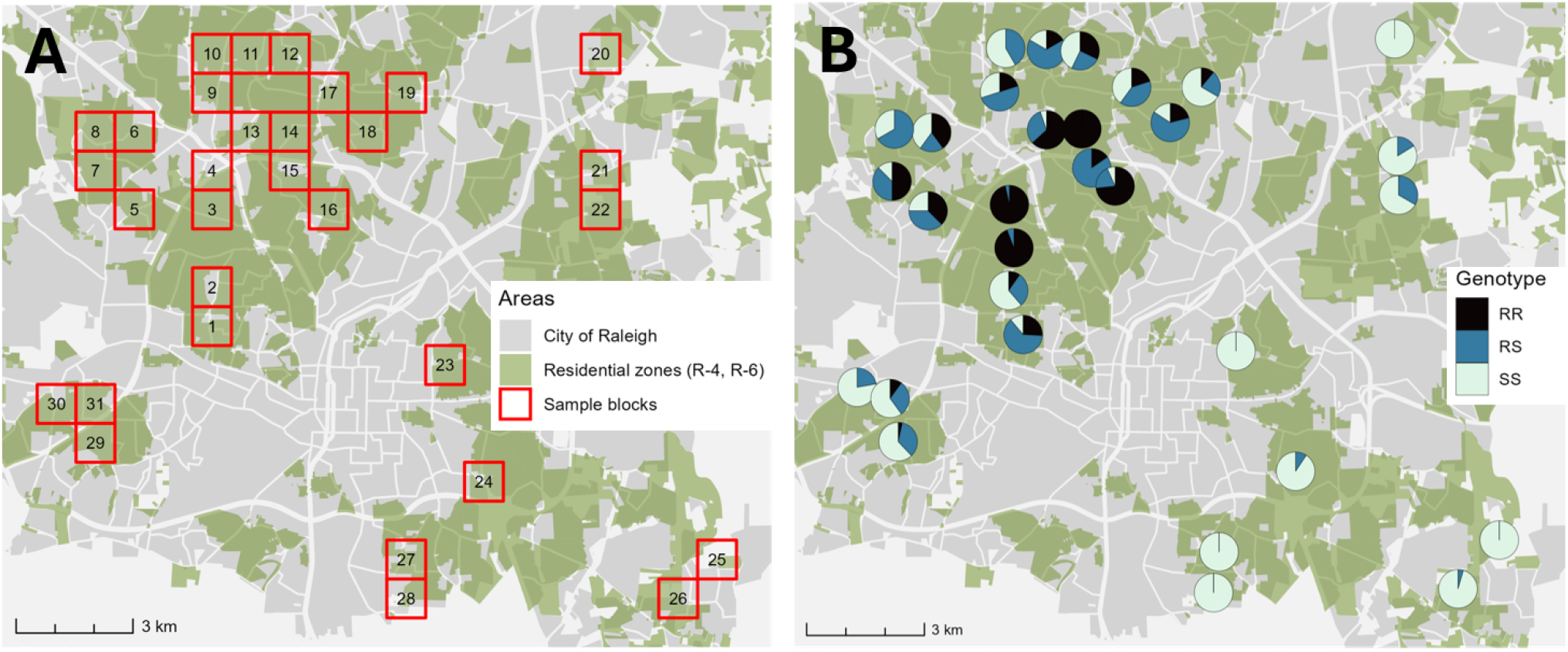
(A) A map of the blocks in Raleigh, NC selected for mosquito sampling. Mosquitoes were collected from sites located near the centroid of each block. Only blocks that have more than 50% of their area zoned as R-4 or R-6 (i.e., four or six homes per acre, respectively) and with centroids composed of single-family homes were selected. (B) Genotype frequencies observed after sequencing. *R* denotes *kdr* mutation that is pyrethroid resistant (F1534S), while *S* denotes the wild-type susceptible allele.

The Moran’s *I* statistic for *kdr* allele frequency was 0.71 (*p <* 0.001; queen contiguity), with the positive value indicating geographic clustering (sample blocks located in closer proximity tend to have more similar *kdr* frequencies). A Moran’s *I* statistic of zero indicates no spatial autocorrelation (*kdr* frequencies are not influenced by neighboring allele frequencies), while a negative statistic indicates a dispersed pattern (sample blocks located in closer proximity have more dissimilar *kdr* frequencies). The explanatory variables demonstrated positive spatial autocorrelation based on Moran’s *I* weighted by inverse distance (*p <* 0.001, *p* = 0.003, and *p* = 0.049 for total value, house age, and abundance, respectively). Local Moran’s *I* analysis showed similar spatial clustering patterns of the response variable and the explanatory variables total property value and house age. After fitting our final regression models with *kdr* allele frequency as the response variable, the model residuals showed no evidence of spatial autocorrelation (*p* = 0.19). This tells us that our models successfully account for the factors leading to spatial clustering of *kdr* allele frequencies [15].

We fit two logistic regression models to the *kdr* data: a main effects-only model, and a model that includes an interaction term between abundance and house age. We considered the latter model because recent work showed that the abundance of *Ae. albopictus* increases with neighborhood age [61, 30], and therefore might affect the frequency of insecticide application. For both models, we used a quasi-binomial distribution to better account for variance in the response [30]. We compared the models using QAICc and found that the main effects-only model was more parsimonious (*Δ*QAICc *≈* 6.2; Table 1) [8]. Prior to removing any outliers, only the intercept (*p <* 0.05) and total property value (*p <* 0.01) were statistically significant. We investigated the influence of individual data points using Cook’s Distance and found Block 2 had a disproportionate effect on the model fit. The explanatory variables recorded for Block 2 were in the extreme regimes of their respective distributions: highest average house age (83 years), a mosquito abundance of 46 per trap effort (90^th^ percentile), and a relatively high property value (81^st^ percentile). We removed Block 2 from our analysis and re-fit the model. When the outlier was omitted, the intercept (*p <* 0.05), house age (*p <* 0.05), and total property value (*p <* 0.001) were significant. Both house age and total property value were found to have a positive association with *kdr* frequency. The McFadden pseudo-*R*^2^ of the fitted model is 66.5% (55.9% with the outlier included). The odds ratios from the GLM fit and the 95% confidence intervals computed from the non-parametric bootstrap are given in Table 1.

**Table 1.** Results of the logistic regression on F1534S frequency, including odds ratio estimates, 95% confidence intervals, and *p*-values of predictors from the most parsimonious model (main effects only, *Δ*QAICc *≈* 6.2). C.I.s and *p*-values are computed from a non-parametric bootstrap with *N* = 10, 000 replicates.

| Coefficients | Odds Ratio | 95% C.I. | $p$-value |
| --- | --- | --- | --- |
| Intercept | 0.638 | [0.385, 1.121] | $< 0.05$ |
| House age | 1.030 | [0.996, 1.067] | $< 0.05$ |
| Abundance | 0.994 | [0.957, 1.025] | 0.303 |
| Total property value | 8.552 | [3.499, 56.354] | $< 0.001$ |

Our statistical model indicates strong evidence that total property value is associated with F1534S mutation frequency. Fig. 2 shows the predicted frequency of F1534S mutations as the total property value varies. (While the statistical fit is given against total property value in Fig. 2, recall that the total property value was log-scaled prior to fitting.) All other predictors are fixed at their mean values. The predicted *kdr* allele frequency is 20% for property values of roughly $350,000, 50% for property values of $665,000, and exceeds 80% for property values of approximately $1.28 million. The intercept is statistically significant (*p <* 0.05), and our model predicts a *kdr* frequency of 39.0% for an average single-family, detached home (i.e., zoned as R-4 or R-6) in Raleigh, NC. The logistic regression model predicts that the odds of *kdr* frequency increase by 3.0% each year after house construction. The predicted *kdr* frequency when all other explanatory variables are fixed at their mean values is 22% at 20 years old, 34% at 40 years, and 48% at 60 years. We found no statistically significant association between mosquito abundance and *kdr* frequency. Figs. 3 and S2 show the predicted *kdr* frequency against house age and abundance, respectively.

**Figure 2.**
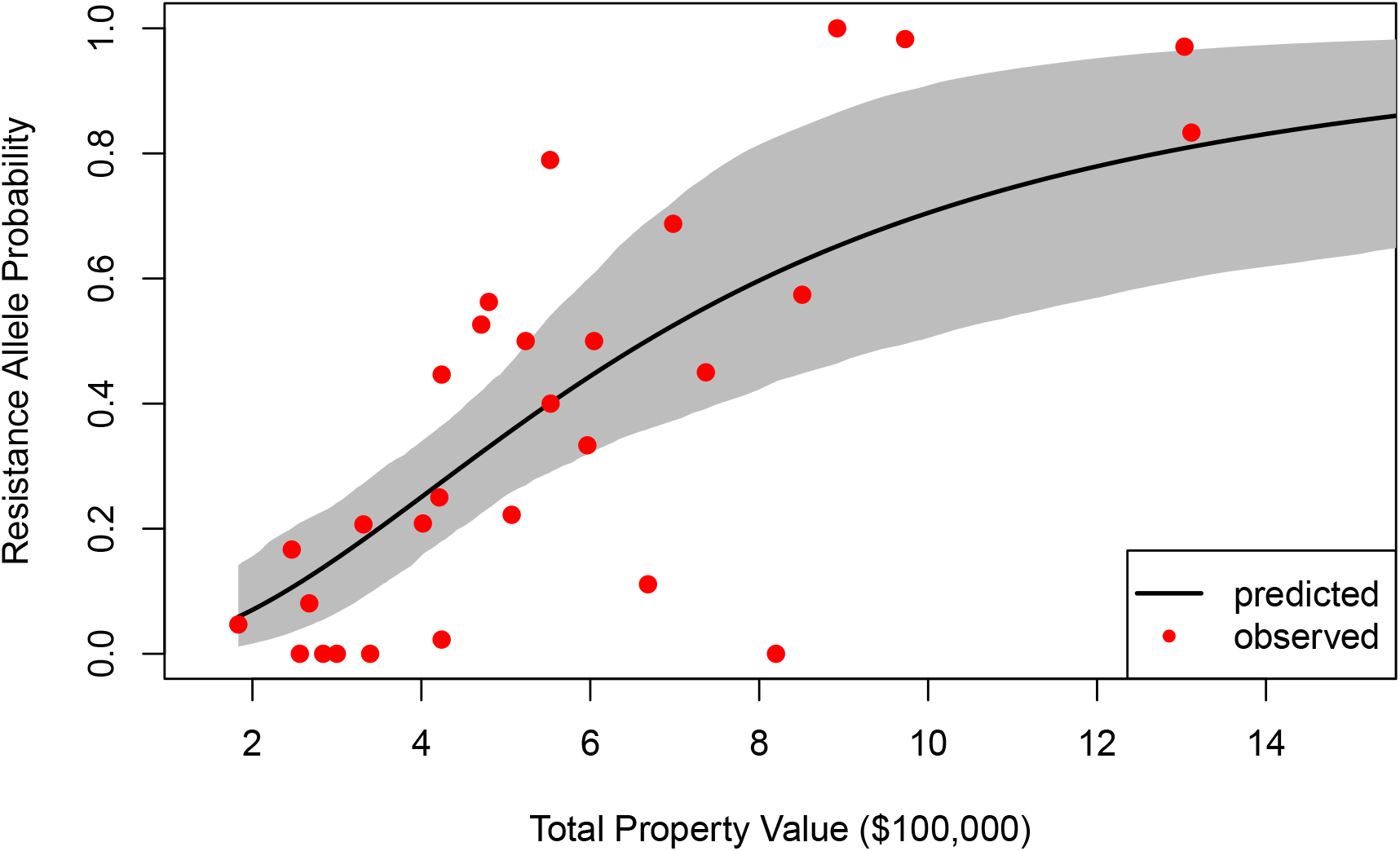
Allele probability of *kdr* mutation against the total property value, in multiples of $100,000. The remaining explanatory variables are fixed at their mean values. The solid black line denotes the expected value of the fitted logistic model; red points denote observed probabilities. A 95% confidence interval of the expected response is computed from 1000 samples of the bootstrapped parameter distributions.

**Figure 3.**
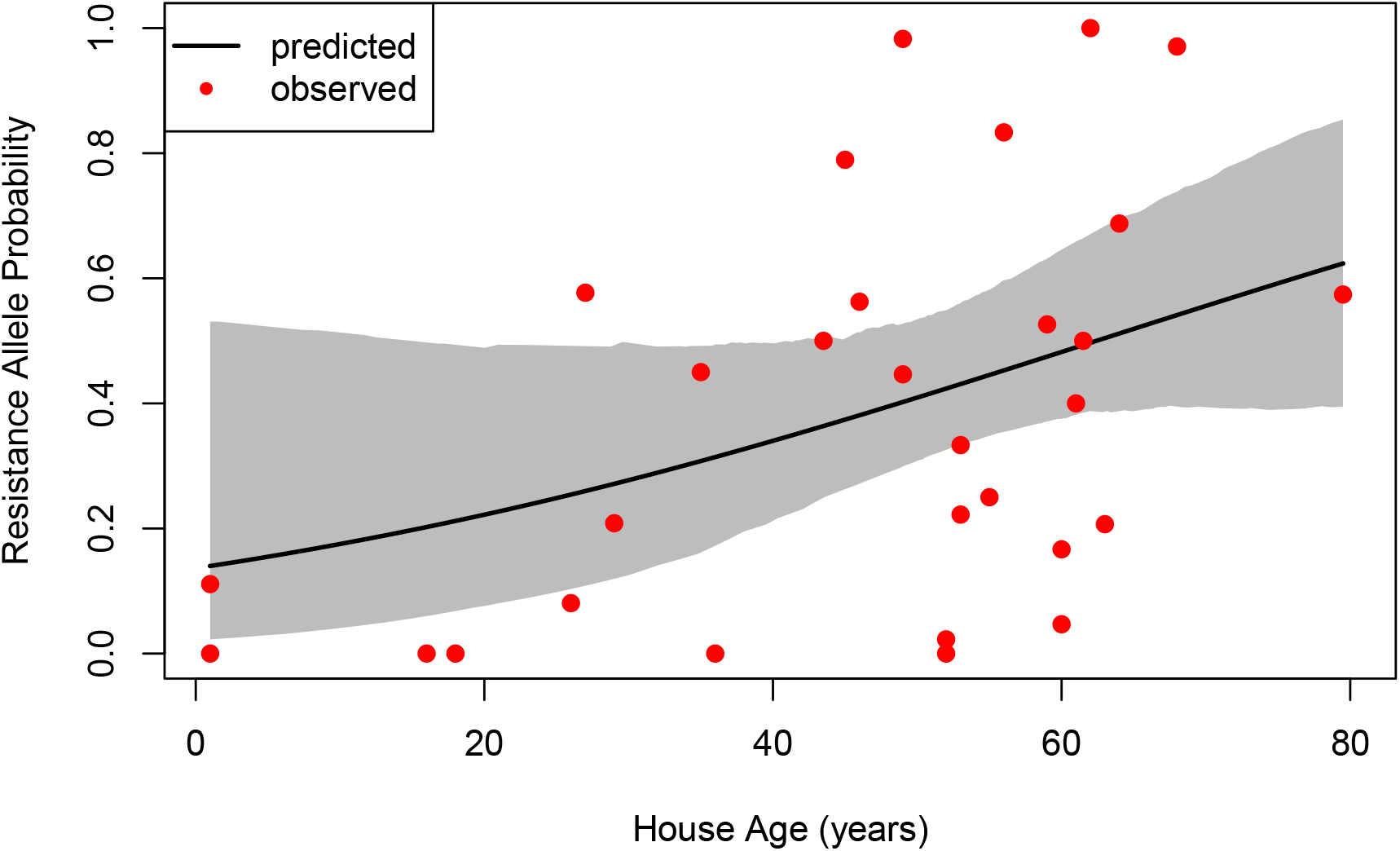
Allele probability of *kdr* mutation against house age. The remaining explanatory variables are fixed at their mean values. The solid black line denotes the expected value of the fitted logistic model; red points denote observed probabilities. A 95% confidence interval of the expected response is computed from 1000 samples of the bootstrapped parameter distributions.

Next, we fit a negative binomial model to the mosquito abundance data using house age and total property value as predictors. We took the response to be the total number of trapped mosquitoes, while the number of 24-hour trapping periods was used as an offset. A chi-squared likelihood ratio test revealed that the dispersion parameter of the negative binomial model was significantly greater than zero (*χ*^2^ = 433; df = 1; *p <* 10^−6^). We used Cook’s Distance to identify potential outliers and removed two sites with the highest mosquito abundance per trap effort. These sites had an abundance per trap effort of 200 and 118—the next highest abundance per trap effort was 66. With the outliers omitted, the intercept (*p <* 10^−6^) and house age (*p <* 0.05) were statistically significant. (With the outliers included, the intercept (*p <* 10^−6^), house age (*p <* 0.01), and total property value (*p <* 0.05) are statistically significant.) The McFadden pseudo-*R*^2^ of the negative binomial model is 14.7% (29.6% with the outliers included). Table 2 shows the results of the negative binomial regression, including incidence rate ratios and their 95% confidence intervals computed from the non-parametric bootstrap.

**Table 2.** Results of the negative binomial regression on mosquito abundance per trap effort, including incidence rate ratios, 95% confidence intervals (C.I.s), and *p*-values of predictors. C.I.s and *p*-values are computed from a non-parametric bootstrap with *N* = 10, 000 replicates.

| Coefficients | Incidence Rate Ratio | 95% C.I. | $p$-value |
| --- | --- | --- | --- |
| Intercept | 10.194 | [6.794, 13.549] | $< 10^{-6}$ |
| House age | 1.022 | [1.000, 1.039] | $< 0.05$ |
| Total property value | 0.949 | [0.600, 2.093] | 0.837 |

Additionally, we fit the negative binomial model using the number of 24-hour trapping periods as a covariate (not shown). Total abundance was not significantly affected by the number of trapping periods (*p* = 0.788), so that trapping results over a single 24-hour trapping period likely reflect overall mosquito abundance at that site. Our statistical findings suggest that the daily abundance of mosquitoes increases by 2.2% each year after construction (see Fig. 4). The expected daily mosquito abundance is approximately 6 for 20-year-old houses, 9 for 40-year-old houses, 13 for 60-year-old-houses, and 21 for 80-year-old houses. These counts are likely lower bounds given the small effective range of mosquito traps (< 15 m. [7]).

**Figure 4.**
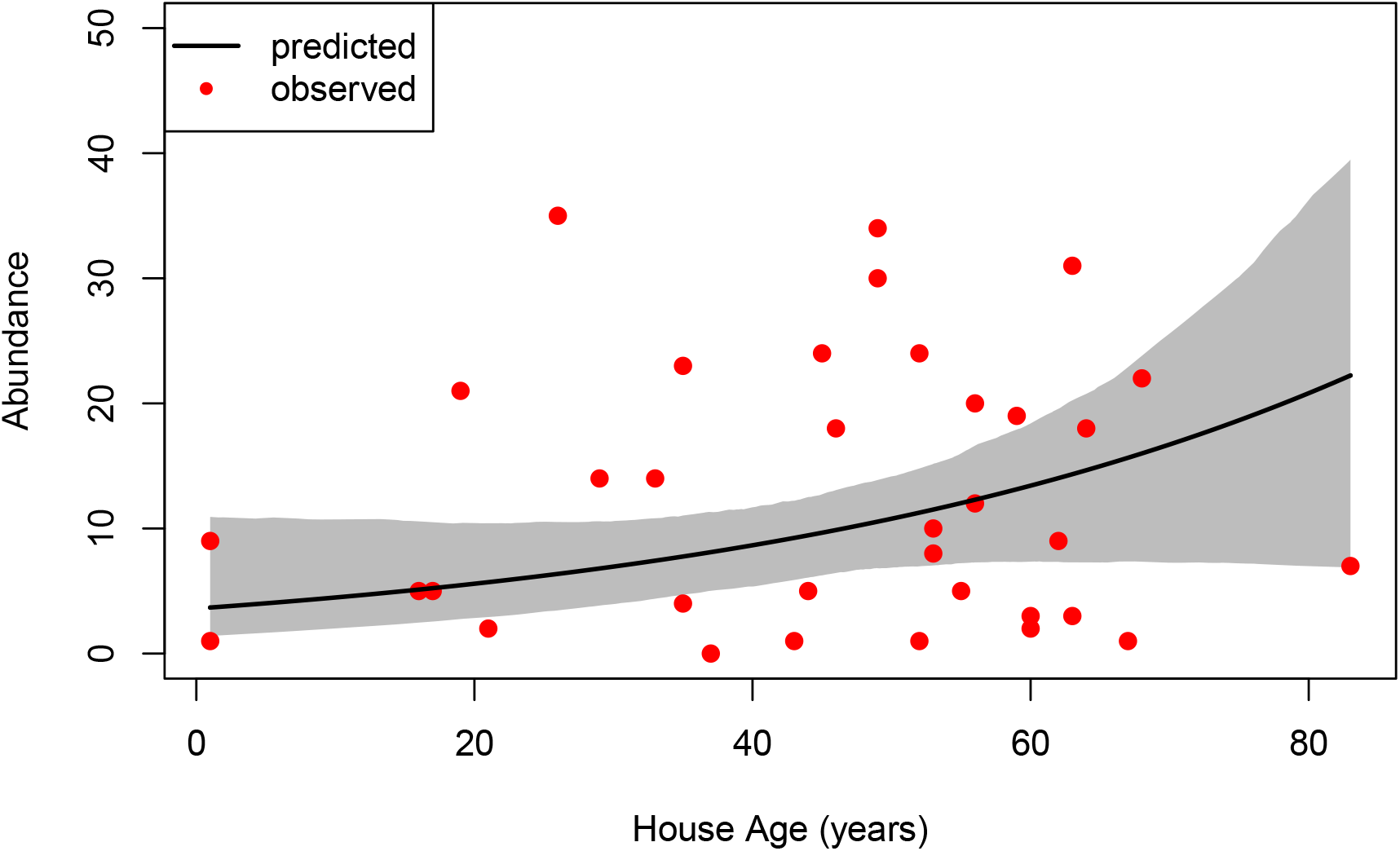
Daily mosquito abundance against house age. Total property value is fixed at its mean value. The solid black line denotes the expected value of the fitted negative binomial model; red points denote observed probabilities. A 95% confidence interval of the expected response is computed from 1000 samples of the bootstrapped parameter distributions.

## 5 Discussion

We found a significant relationship between wealth, measured by total property value, and the frequency of *kdr* mutations in *Ae. albopictus* populations in Raleigh, NC. We speculate that the spraying of chemical insecticides is more frequent in affluent neighborhoods, as insecticide exposure drives an increase in *kdr* mutations in *Ae. albopictus* and other mosquito species [4, 39, 41, 59, 72]. Current research supports this speculation. In their survey, Richards *et al*. (2017) found that North Carolina residents in moderate to high socioeconomic categories applied insecticides to their properties more often than those from low socioeconomic categories [55]. Elsewhere, socioeconomic status has been shown to correlate with more frequent use of vector control measures, including insecticide spraying and bed nets [19, 34]. Dickinson et al. (2022) found that a resident’s willingness to pay for mosquito control increased with income [14]. The link between socioeconomic status and pyrethroid resistance has previously been explored in the related species Ae. aegypti: Fay *et al*. (2023) found a higher frequency of pyrethroid resistance in *Ae. aegypti* in two wealthier neighborhoods compared to two poorer neighborhoods in Argentina [20]. In future work, we will examine the explicit connection between wealth, insecticide exposure, and the evolution of resistance.

The resistance frequencies we study here are part of a historical trend over multiple generations [5]. This is supported by the high frequency of resistant homozygotes (R/R; 25.8%), which take longer to appear following the emergence of *kdr* alleles [44, 64]. Since the last survey in 2020 for *kdr* in *Ae. albopictus* in Raleigh, NC [1], the frequency of F1534S has increased considerably. Only 4.3% (12 of 278) of *Ae. albopictus* screened by Abernathy *et al*. (2022) had *kdr* mutations; these samples were all taken from two of nine residential sites [1]. In the present study, we found that 52.1% (279 of 535) of *Ae. albopictus* mosquitoes spanning 26 of 31 residential blocks had a *kdr* mutation. We show elsewhere that these findings are part of a historical trend of increasing *kdr* mutation range and frequency, and calculate a selection coefficient consistent with positive selection [5]. This is not unusual: similar work has found rapid spread of *kdr* mutations in mosquitoes in response to pesticide application [4, 62]. Su *et al*. (2018) found that mortality to permethrin (a Type I pyrethroid) fell by 40% in one population of *Ae. albopictus* after only a year [62]. Rapid spread of *kdr* mutations in response to increased pesticide use has been observed in other mosquito species, including *Ae. aegypti* [4], *Anopheles gambiae* [41, 51], and *Culex pipiens* [59].

Our data provide evidence for an association between *kdr* frequency and wealth. We speculate this association is due to a greater frequency of insecticide spraying in affluent neighborhoods [55], demonstrated by a higher willingness to pay for mosquito control [14]. We also found that house age was positively associated with F1534S frequency and mosquito abundance. This association could be explained by older neighborhoods supporting greater densities of *Ae. albopictus* through increased oviposition sites [61] and therefore leading to more frequent insecticide use. Our logistic regression model predicts a positive *kdr* frequency for an average single-family detached home in Raleigh, NC, as the intercept was statistically significant. We did not find evidence of an association between mosquito abundance and *kdr* mutation frequency. Indeed, how adult mosquito density and *kdr* frequency could be linked is complex and likely depends on the timing of control. For example, high densities could indicate a resistant and thriving population or a population for which there is no vector control. Furthermore, we did not find a statistically significant association between total property value and *Ae. albopictus* abundance. Indeed, evidence for a general association between socioeconomic status and mosquito abundance is mixed and appears to be region-specific [37, 60, 73, 74].

We found that *kdr* frequency was correlated in space. The spatial clustering in *kdr* allele frequency is likely due to clustering in the explanatory variables, based on a Moran’s *I* analysis. Indeed, the highest frequencies of *kdr* were located in older neighborhoods with high property values in the suburbs of northwest Raleigh. Our model residuals were spatially uncorrelated, indicating that the explanatory variables successfully accounted for the spatial autocorrelation in *kdr* frequency. This fact, combined with the overlap between spatial patterns of property value and *kdr* frequency, further supports our hypothesis.

Gene flow acts to genetically homogenize populations. In our case, however, we expect gene flow to be limited by low organism dispersal [42, 63]. When local selection is strong and gene flow is weak, a mosaic allele frequency occurs characterized by high variance [26]. Mosquito control in Raleigh is performed at the household-level by residents and private contractors, so that the application of chemical insecticides is likely heterogeneous. One recent survey of NC residents found that 31% of respondents apply insecticides to their properties to control mosquitoes, although only 5% engaged in commercial backyard control [55]. The combination of low gene flow and heterogeneous insecticide spraying could explain the high spatial variance of F1534S frequency we observed, even when comparing adjacent blocks. The high frequency of F1534S homozygotes could indicate persistent (i.e., year-to-year) insecticide application. Variance in resistance patterns at local scales has been found elsewhere for *Ae. albopictus* [22], as well as for the related species, *Ae. aegypti* [4, 26, 47].

There are several ways our study can be improved. First, our sampling was random and did not uniformly cover the range of conditions spanned by the explanatory variables. Informed sampling algorithms, such as that proposed in [73], would have extended the range of conditions over which we could make inference about the association between the response and explanatory variables. Nonetheless, the range of conditions explored in the present study is broadly representative of suburban landscapes characterized by residential single-family homes. We have been careful to reserve inference for only those areas of parameter space represented by our data. Furthermore, we used a non-parametric bootstrap to compute confidence intervals and conduct hypothesis tests given our small sample sizes. The non-parametric bootstrap assumes that our sample is representative of our population (i.e., single-family homes in Raleigh, NC), but we thought this to be a safer assumption than those made by alternative methods relying on asymptotic theory. We opted for a simple and efficient non-parametric bootstrap while improved methods have been proposed [13]. Nonetheless, we expect that the statistically significant associations we found would be unchanged if we had used more advanced techniques. We found that house age was barely statistically significant as a predictor of F1534S frequency (*p* = 0.0471). A statistical analysis with a larger and more diverse sample would be useful to increase our understanding of the relationship between insecticide resistance and neighborhood age.

Future work can most immediately investigate our assumed relationship between resident wealth and insecticide spraying frequency. An analysis exploring the population genetic structure of our samples would help ascertain the origin of *kdr* in Raleigh, NC mosquitoes, and clarify whether the present extent of pyrethroid resistance is due to *de novo* mutations or via dispersing mutants. Finally, more studies of the association between *kdr* frequency in *Ae. albopictus* and socioeconomic predictors are useful to guide and inform vector control efforts in the future.

In this work, our data show strong evidence that *kdr* frequency in *Ae. albopictus* is positively correlated with wealth, measured by total property value. The predicted *kdr* frequency is 20% for property values of roughly $350,000 but reaches 70% for property values exceeding $1,000,000, assuming all other explanatory values are fixed at their average values. It is likely that this association is due to higher frequencies of insecticide application in wealthier neighborhoods. However, we are yet to demonstrate a connection between wealth and spraying frequency. We also found that house age is positively associated with *kdr* frequency and *Ae. albopictus* abundance, suggesting that spraying is more frequent in older communities where mosquito densities are higher. By investigating the relationship between wealth and *kdr* frequency of *Ae. albopictus*, these findings can guide vector control efforts accordingly.

## Supplementary Information

**Figure S1.**
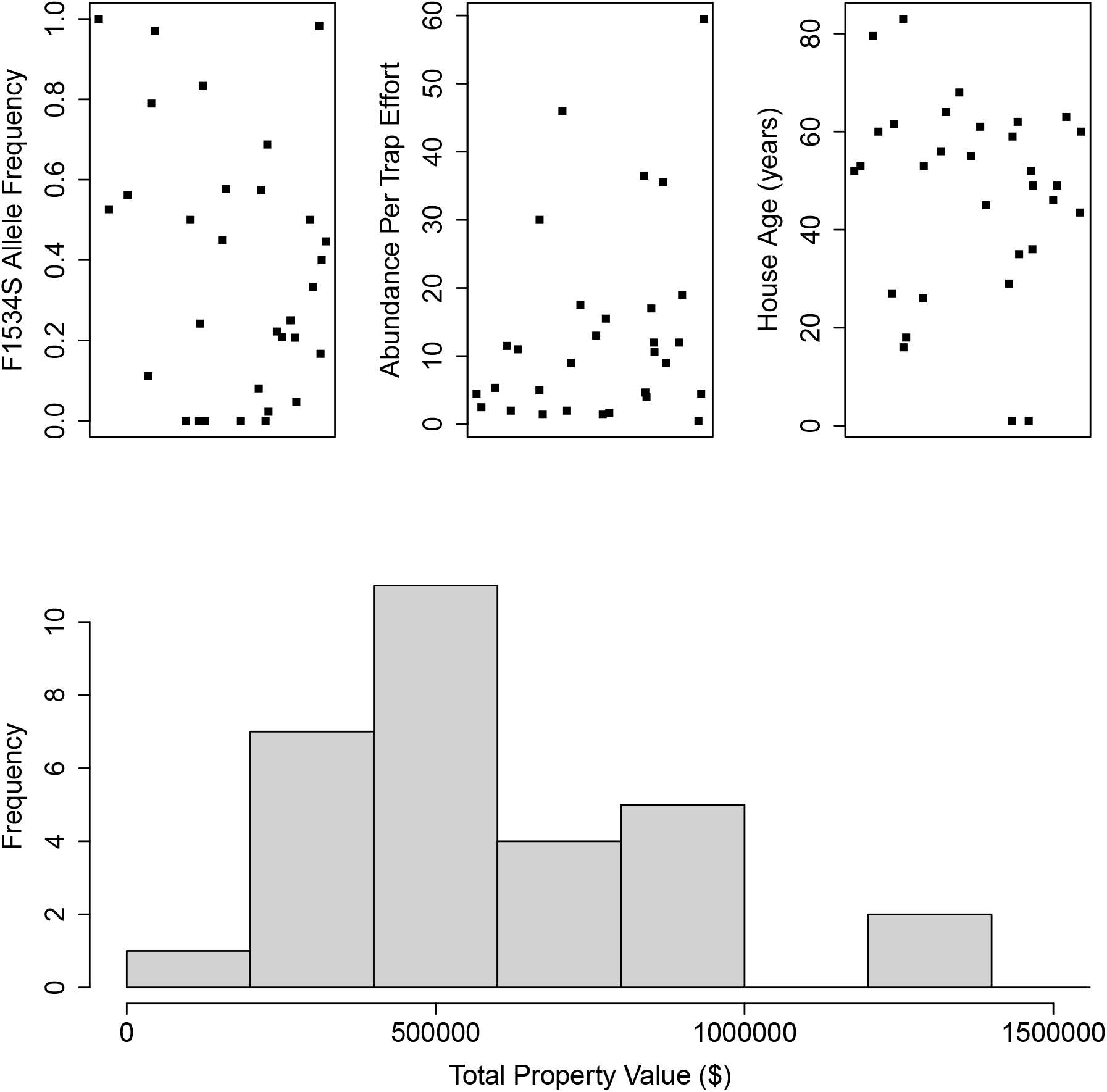
Plots of the data for each explanatory variable used in the statistical analysis. For each block, we recorded average abundance per trap effort, total property value (log-scaled), and house age (relative to 2023). The data here are not scaled for ease of interpretability. Note that total property value is the sum of home and lot value; a histogram of home value is also provided. A single point was omitted from the scatter plot of abundance as the recorded mean value for this block was 200 mosquitoes per trap effort.

**Figure S2.**
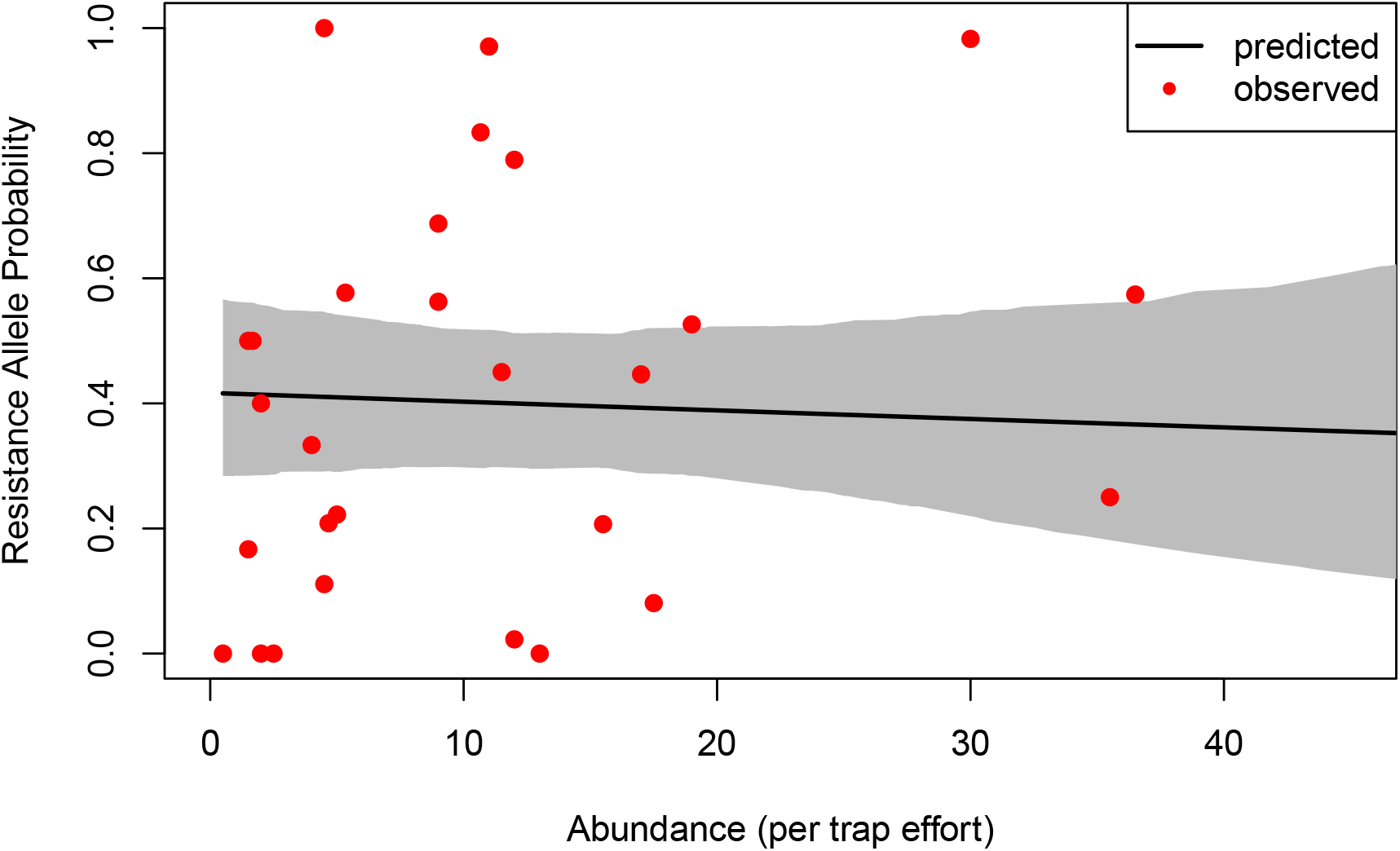
Allele probability of *kdr* mutation against abundance per trap effort. The remaining explanatory variables are fixed at their mean values. The solid black line denotes the expected value of the fitted logistic model; red points denote observed probabilities. A 95% confidence interval of the expected response is computed from 1000 samples of the bootstrapped parameter distributions.

